# Crossmodal congruency effect scores decrease with repeat test exposure

**DOI:** 10.1101/186825

**Authors:** Satinder Gill, Daniel Blustein, Adam Wilson, Jon Sensinger

**Affiliations:** Institute of Biomedical Engineering, University of New Brunswick, Fredericton, NB, Canada; Department of Electrical and Computer Engineering, University of New Brunswick, Fredericton, NB, Canada

**Keywords:** Crossmodal congruency, Learning effect, multisensory integration, reaction time

## Abstract

The incorporation of feedback into a person’s body schema is well established. The crossmodal congruency effect (CCE) task is used to objectively quantify incorporation without being susceptible to experimenter biases. This visual-tactile interference task is used to calculate the CCE score as a difference in response time for incongruent and congruent trials. Here we show that this metric is susceptible to a learning effect that causes attenuation of the CCE score due to repeated task exposure sessions. We demonstrate that this learning effect is persistent, even after a 6 month hiatus in testing. Two mitigation strategies are proposed: 1. Only use CCE scores that are taken after learning has stabilized, or 2. Use a modified CCE protocol that decreases the task exposure time. We show that the modified and shortened CCE protocol, which may be required to meet time or logistical constraints in laboratory or clinical settings, reduced the impact of the learning effect on CCE results. Importantly, the CCE scores from the modified protocol were not significantly more variable than results obtained with the original protocol. This study highlights the importance of considering exposure time to the CCE task when designing experiments and suggests two mitigation strategies to improve the utility of this psychophysical assessment.

## Introduction

There is an increasing interest concerning the multisensory representation of space in humans due to converging findings from a number of different disciplines. The crossmodal congruency task is a visual-tactile interference task that has been used to investigate multisensory representation of space in humans (Spence, Nicholls, Gillespie & Driver 1998; Spence, Pavani & Driver 2004; Spence, Pavani, Maravita & Holmes 2004), including those with brain damage (Spence, Kingstone, Shore & Gazzaniga 2001). Investigation of crossmodal selective attention has been used to demonstrate the detrimental effects of age on the ability to ignore irrelevant sensory information when attending to relevant sensory information (Poliakoff, Ashworth, Lowe & Spence 2006). Other studies have investigated changes in the representation of peripersonal space that are elicited by the prolonged use of hand-held tools (Holmes, Calvert & Spence 2007; Holmes & Spence 2004; Maravita, Spence, Kennett & Driver 2002). Further studies used this task to extend findings regarding physical and pointing tools to virtual robotic tools using techniques from haptics and virtual reality (Sengül et al., 2012; Sengül et al., 2013). Recently it has been used in conjunction with the rubber hand illusion paradigm to investigate realistic rubber hand incorporation into body schema (Zopf, Savage & Williams 2010; Zopf, Savage & Williams 2013). The crossmodal congruency effect (CCE) assessment has become a well-established quantitative metric of feedback incorporation that extends to a variety of applications.

Typically, CCE study consists of a participant holding two foam blocks in either hand with vibrotactile targets and visual distractors embedded in the top and bottom of each foam block. A trial consists of a random and independent presentation of a single vibrotactile target paired with a single visual distractor from any of the four possible pairs of locations. Participants are instructed to make a speeded response regarding the elevation of the vibrotactile target (i.e. “up”, at the index finger verses “down”, at the thumb), while simultaneously ignoring visual distractors (Spence et al., 2001). Participants are typically slower at discriminating the elevation of vibrotactile targets when visual distractors are presented from an incongruent elevation (i.e. when vibrotactile target and visual distractors are presented from different elevations) as compared to a congruent elevation. The CCE score is calculated as a difference in the reaction time for congruent and incongruent trials and is used as a quantitative performance metric for multisensory representation of space. It has been demonstrated that CCE scores are typically higher when the spatial separation between visual distractor and vibrotactile targets is low (e.g. both locations on same block vs. locations on two different blocks held in different hands) (Spence, Pavani & Driver 2004; Maravita, Spence & Driver 2003). CCE is a simple stereotypical behavioral task that provides a robust performance metric of feedback incorporation.

Despite the widespread use of the crossmodal congruency task in a variety of experimental paradigms, important questions related to task exposure times remain unanswered. For example, in a majority of the studies the experimental protocol consists of multiple blocks of trials with experimental sessions lasting up to approximately 60 minutes. These time-intensive experimental sessions might result in extended learning of the task and adversely affect a participant’s CCE score. Also, there is no specific mention of a participant selection criterion based on previous knowledge of the crossmodal congruency task. To our knowledge, there exist no studies to date that have investigated modulation of CCE score due to repeated task exposures. If a task learning effect exists for the CCE, results from previous studies that ignored such an effect may be misinterpreted. Participants tested with previous CCE experience would be expected to have lower CCE scores than participants naïve to the assessment. A learning effect that has been ignored by the field may render inaccurate our current understanding of multisensory space representation in humans. In this study we present evidence that CCE scores decrease with repeated task exposure sessions and that this learning effect is persistent over time. We propose two mitigation strategies, including a shortened CCE protocol that we compare to the established protocol. Best practices for future CCE score use are suggested.

## Materials and Methods

### CCE test implementation

During CCE testing, participants sat comfortably on a chair in front of a table in a dimly illuminated room. They wore over-ear noise-canceling headphones playing Brownian noise to mask background noise. Identical vibrotactile stimulation motors were placed on the thumb and index fingertips of the participant’s right hand (see the Materials and Apparatus section for hardware details). Distractor light emitting diodes (LEDs) were placed on the thumb and index finger of the participant’s right hand while a fixation LED was centered between the distractor LEDs using a plastic mounting strip. Fig 1 shows the system setup used for this experiment.

**Fig 1.**
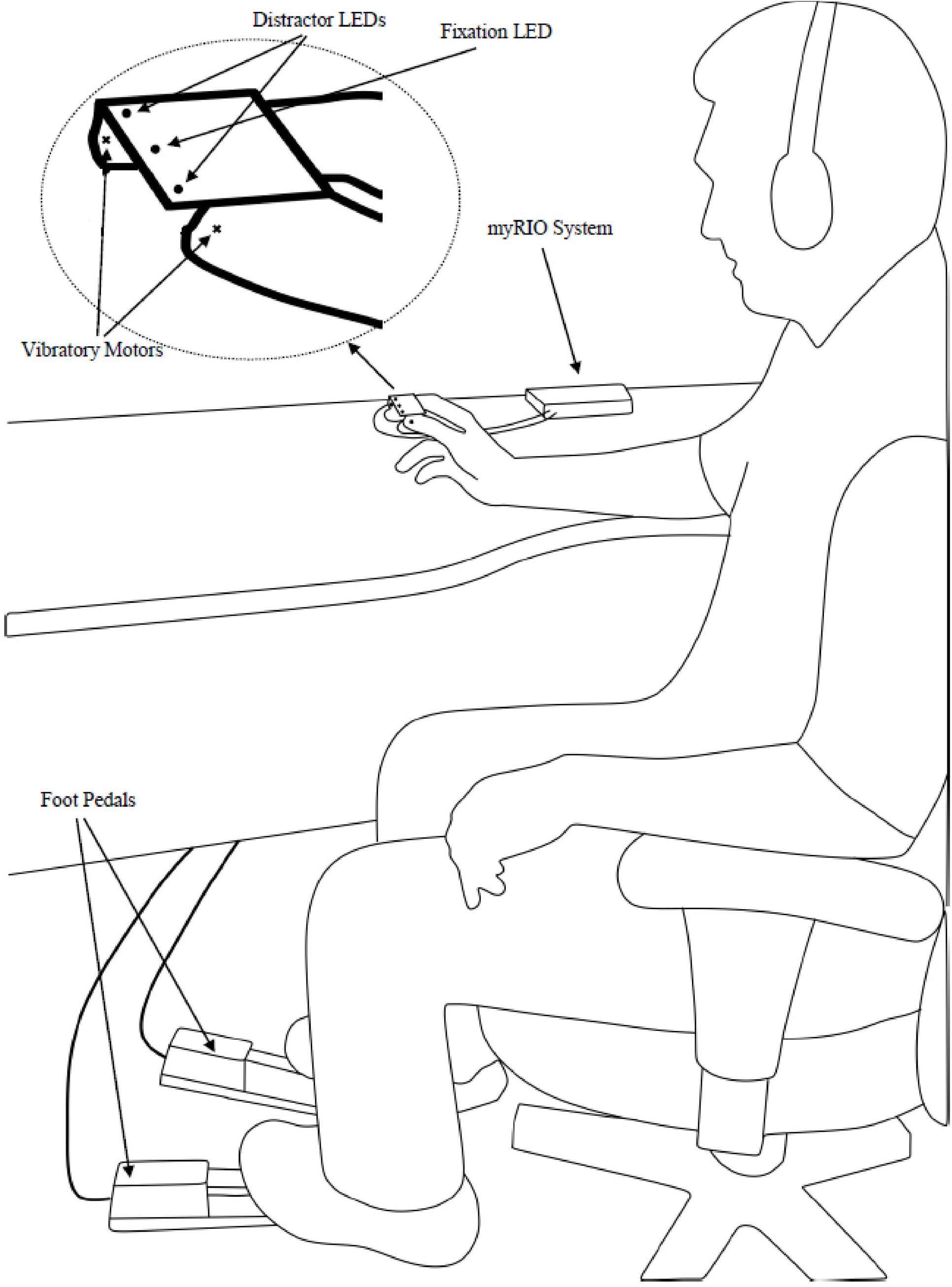
Overall system setup. Subjects make speeded responses with foot pedals to vibratory feedback presented on the index finger or thumb. Subject is asked to focus on the fixation LED while ignoring the distractor LEDs. See text for details.

At the beginning of the CCE assessment, participants were instructed to make a speeded response to vibrotactile targets that were presented randomly to either the thumb or index fingertip. Participants responded by pressing the left or right foot pedals to indicate stimulation of the thumb or index finger, respectively. Participants were explicitly instructed to respond as fast as possible, while making as few errors as possible. Participants were also informed that visual distractors would be presented simultaneously with the vibrotactile targets but that they were completely irrelevant to the vibrotactile target discrimination task. They were specifically instructed to ignore visual distractors by keeping their eyes open and fixating their vision on the central fixation LED. The fixation LED was presented 1000ms before the vibrotactile targets and visual distractors were simultaneously presented for 250ms.

The CCE test session was preceded by a practice session consisting of three blocks of 10 trials each. The first block of the practice session consisted of presenting the fixation LED and vibrotactile targets only (i.e. no visual distractors were presented) so that participants could familiarize themselves with the vibrotactile target discrimination. The visual distractors were presented along with vibrotactile targets in the following two blocks of the practice session. This was followed by the experimental session during which 64 trials were presented to each participant during each block of trials. Either four or eight blocks of trials were presented depending upon the particular experiment. Each trial ended when the participant responded by pressing one of the foot pedals or no response was made within 1500ms of target onset.

### Participants

A total of 30 healthy participants were recruited from the local community (23 male, 7 female; aged 19 – 57 years, mean ± SD = 27.5 ± 7.8 years; 26 right hand dominant, 1 left hand dominant, 3 reported no particular hand dominance). All participants had normal or corrected to normal vision, no disorder of touch, were able to use both foot pedals, and were naïve to the CCE task. Participants were informed about the general purpose of the research and were given the opportunity to ask questions but they were not aware of the specific goals of the research. Experiments were conducted under human ethics approval at the University of New Brunswick (Fredericton, NB, Canada) and the U.S. Department of the Navy’s Human Research Protection Program. Written informed consent was obtained from each participant before conducting experiments and no compensation was provided to participants.

### Materials and apparatus

The experimental test platform was designed around the National Instruments (NI) myRIO embedded hardware system to achieve millisecond timing accuracy. The three digital outputs of the myRIO system were used to drive one 3mm green LED, termed the fixation LED, and two 3mm green LEDs named distractor LEDs. Two 308-107 Pico Vibe 8mm vibratory motors from Precision Microdrives were used as the vibrotactile targets for the thumb and index finger. The analog outputs of the myRIO system were used to drive STMicroelectronics L272 power operational amplifiers that drove the vibratory motors. The amplifiers were necessary to provide sufficient driving current to the vibratory motors. Two OnStage KSP100 keyboard sustain pedals were interfaced to the myRIO system through digital inputs to measure a participant’s speeded responses to the vibrotactile targets. NI LabVIEW was used to develop the firmware for the myRIO system to randomly activate the visual distractors and vibrotactile targets at a fixed interval and measure the speeded response time. A desktop NI LabVIEW graphical user interface (GUI) was designed to interact with the myRIO embedded system, allowing the experimenter to set various experimental parameters such as vibratory stimulus amplitude, the number of trials and to specify the filename to record the timing results of speeded responses.

### Experiment design

Three experiments were run to examine CCE learning effects. In Experiment 1 we assessed the learning effect over five exposures to the CCE task. Twelve subjects each completed five consecutive days of CCE testing using the standard 8-block protocol (Spence, Pavani & Driver 2004). The number of sessions and subjects was determined using a power analysis using preliminary CCE results from eight pilot subjects (see Appendix). Subjects were naïve to the CCE and testing was run at about the same time each day. Before each test session, generalized reaction time was measured using a ruler drop task. Subjects held their thumb and first finger 4 cm apart with a vertically-aligned ruler held by the experimenter set so the bottom of the ruler was just within the grasp of the subject. The subject was asked to grasp the ruler as quickly as possible after they noticed it falling. The researcher dropped the ruler at an unspecified and variable time and recorded the distance fallen as a proxy for reaction time. The ‘reaction distance’ was calculated as the average result of three ruler drop trials. In Experiment 2, the persistence of the CCE learning effect was assessed. Eight subjects from Experiment 1 returned for an additional CCE test session after more than six months with no CCE exposure. For both Experiments 1 and 2, all CCE sessions consisted of eight blocks of 64 trials each. Each block lasted for approximately four minutes and a break period of two minutes was provided between consecutive blocks.

In Experiment 3, we examined the effect of reducing the number of CCE blocks per exposure from eight to four. 18 subjects were tested across two exposures on consecutive days; half of the subjects (n=9) were randomly assigned to test with four blocks and the other nine subjects tested with eight blocks for each session.

### Analysis

Practice session trials were not analyzed. To calculate the CCE score from the test trials, we first discarded trials with an incorrect response, trials with a premature response (i.e. reaction time less than 200ms), and trials with a delayed response (i.e. reaction time greater than 1500ms) as these conditions most likely occurred due to lapses in attention (Spence, Pavani & Driver 2004; Sengül et al., 2012). The remaining trials in each block were used to calculate the mean congruent and mean incongruent reaction times. The CCE score for each block was calculated by taking the difference between mean incongruent and congruent reaction times. The mean of the block CCE scores was used to calculate the CCE score for each participant for a particular exposure session. Selection error rates were used as a separate metric of analysis.

In Experiment 1, attenuation across sessions was evaluated using a one-way repeated-measures ANOVA. For Experiment 2, we ran an intraclass correlation coefficient analysis [two-way mixed model with single measurements] and calculated the standard error of measurement 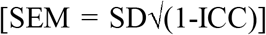 to assess the test-retest reliability between the 6+ month follow-up visit and the Day 5 visit (Weir 2005). Experiment 3 data were analyzed using paired t-tests within the data collected using each protocol. All data are available on Dryad at doi:10.5061/dryad.150v8g3.

## Results

### Experiment 1: CCE score decreases over repeated exposures

We measured the CCE score of 12 subjects across five sessions using the conventional CCE method of 8-blocks. We observed a significant effect of exposure number on CCE score determined with a repeated-measures ANOVA with Greenhouse-Geisser correction (Fig 2) (F(2.00,21.95)=6.93, p = .005, partial-eta=.39). Tests of within-subjects polynomial contrasts over exposure number indicated a significant linear trend (F(1,11) = 8.63, p=.013), a significant 4^th^ order trend (F(1,11) = 5.14, p=.044), and a quadratic trend (F(1,11) = 4.83, p=.05). The statistically-supported trends match the observed initial decrease in CCE score and subsequent stabilization, indicating a task learning effect (Fig 2).

**Fig 2.**
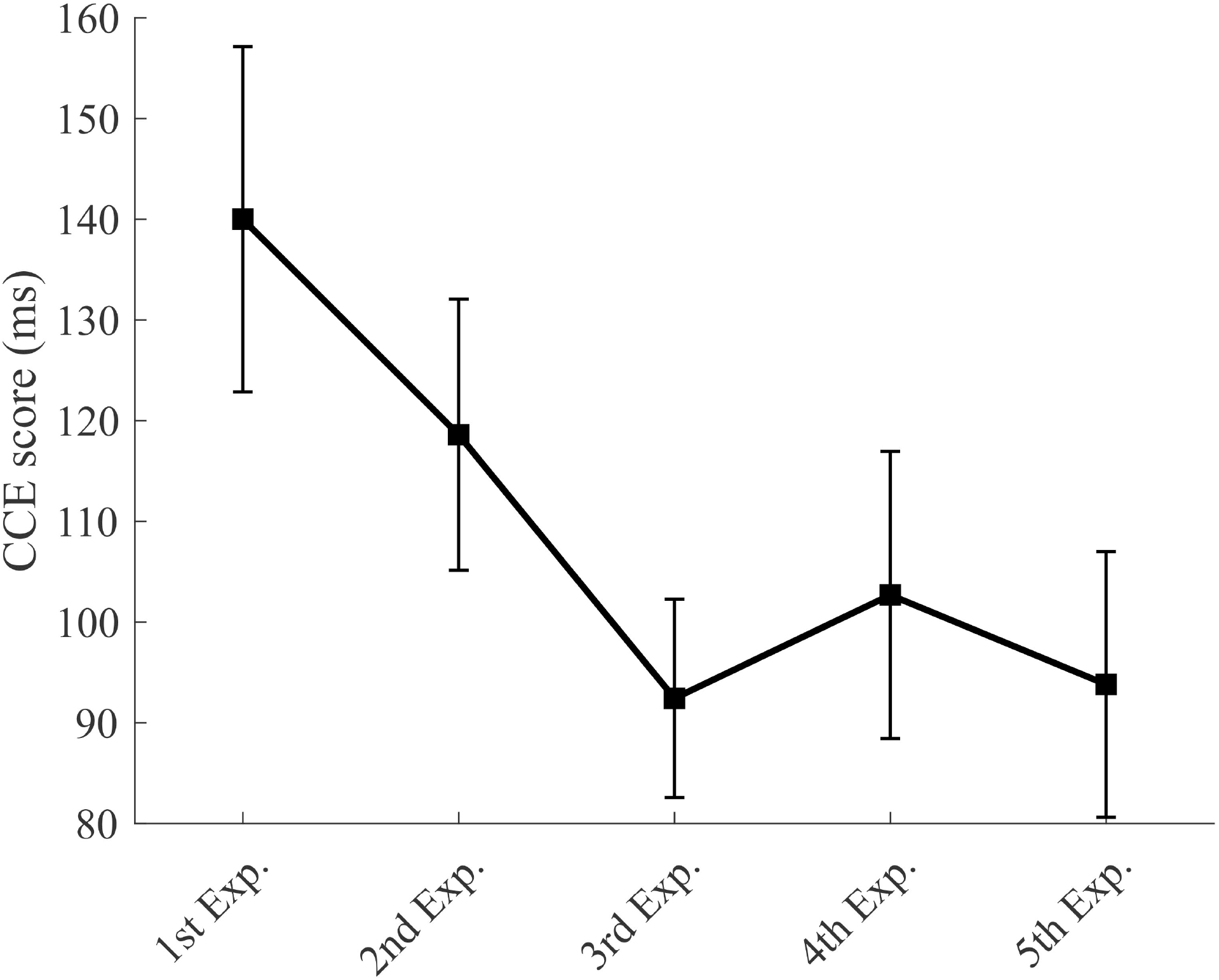
CCE score decreases with task exposure. Each data point represents the mean CCE score for 12 subjects. Error bars show standard error.

To verify that the change in CCE score was due to a learning effect and not attributed to other interacting factors, we analyzed the CCE selection error rates, generalized reaction times measured with a ruler grasp task, and overall reaction times for congruent and incongruent stimuli. We found no significant effect of exposure number on CCE error rate (Fig 3a), as indicated by a repeated-measures ANOVA with Greenhouse-Geisser correction (F(1.15,12.66)=0.53, p = .51). Repeated-measures ANOVA with Greenhouse-Geisser correction indicated a significant effect of exposure on both incongruent reaction time (F(2.16,23.78)=13.57, p <.01) and congruent reaction time (F(1.46,16.07)=8.89, p<.01) (Fig. 3b). To summarize, the only metrics presenting similar exposure-dependent decreases as the CCE scores (Fig 2) were the congruent and incongruent reaction times (Fig 3b). We found no significant effect of exposure number on generalized reaction time as measured by a ruler drop fall-to-grasp distance, as indicated by a repeated-measures ANOVA with Greenhouse-Geisser correction (F(1.463,14.63)=1.269, p = .29).

**Fig 3.**
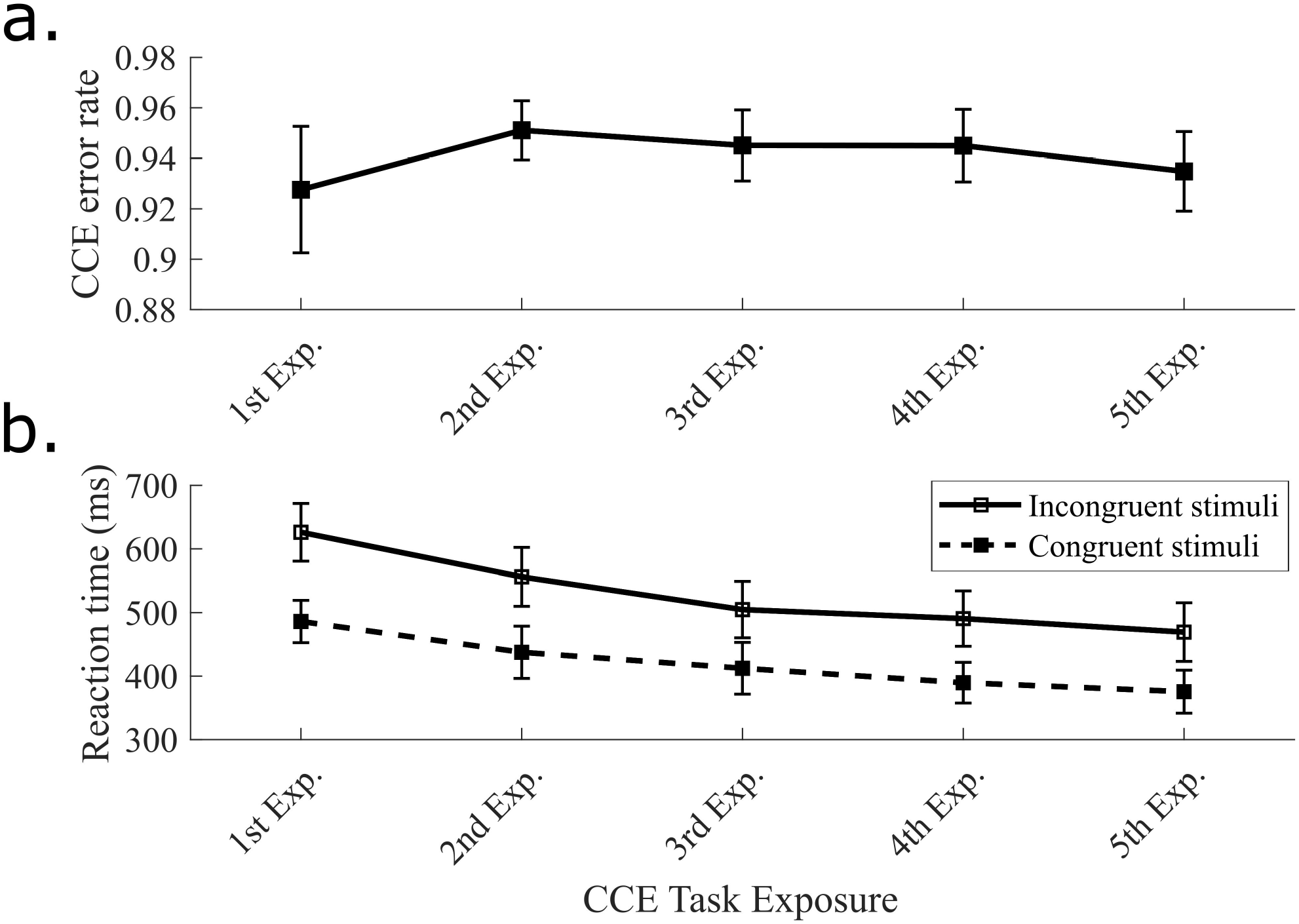
Decreasing CCE score trend matched in isolated reaction times but not in CCE error rates. a. Plot of selection error rate over repeated exposures. An error occurred when the wrong pedal was pressed (e.g. vibration stimulus on the thumb resulted in subject pressing the pedal indicating an index finger stimulus). b. Congruent and incongruent reaction times plotted independently.

### Experiment 2: CCE learning effect persists over time

After observing a statistically significant CCE learning effect, observed in both incongruent and congruent reaction times but not in error rates and generalized reaction times, we wanted to see if the learning effect persisted over longer time periods. Eight subjects who completed Experiment 1 were re-tested between 6 and 7 months after their initial testing. Revisit CCE scores were similar, and even slightly lower, than the CCE scores measured on Exposure 5, about 6 months prior (Fig 4). Similarity between Exposure 5 and the re-visit session was indicated by a high intraclass correlation coefficient (ICC(3,1) = .71) and a low standard error of measurement (SEM = 21.6ms) compared to baseline reliability data calculated by comparing exposure 1 and re-visit results (ICC(3,1) = .46; SEM = 45.7ms).

**Fig 4.**
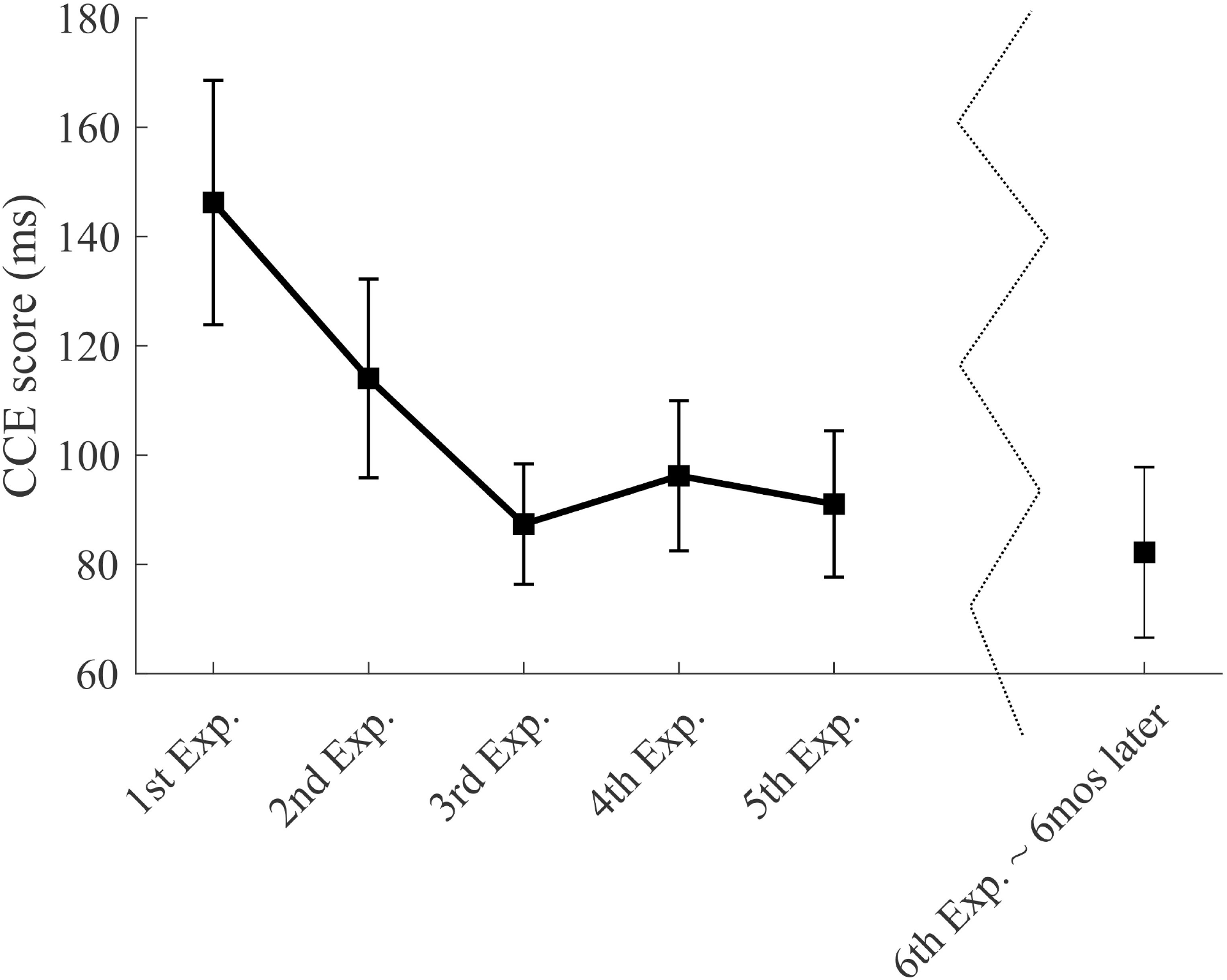
Persistent CCE learning effect after 6-month follow-up testing. Eight subjects from Experiment 1 returned for a sixth visit about 6 months after their initial visits. Data shown are means and standard error for those eight subjects.

### Experiment 3: Modified CCE protocol reduces learning effect

We next sought to explore mitigation strategies to diminish the impact of the CCE learning effect on research results. The persistence of the learning effect suggests that CCE scores should only be considered after task learning has stabilized. However, in certain research or clinical settings with subject access or time constraints, extended testing may be impractical. In Experiment 3 we tested a modified CCE protocol designed to reduce task exposure to determine if the shortened testing could: 1. Produce valid CCE results; and 2. Mitigate the learning effect.

The modified protocol reduced the duration of task exposure in each session from eight blocks to four blocks. The 4-block protocol reduced the drop in CCE score from first exposure to second exposure (Fig 5). For the 8-block protocol, mean CCE score dropped by 14.7ms from the first exposure to the second, compared to a 7.6ms drop for the 4-block protocol. Variability in the second session was only slightly higher for the 4-block protocol (SD = 43.8ms) compared to the 8-block protocol (SD = 39.0ms). The variability difference was more pronounced when analyzing the first session results [4-block SD =64.9ms; 8-block SD = 47.5ms]. None of these differences were statistically significant (paired t-tests, p>.05).

**Fig 5.**
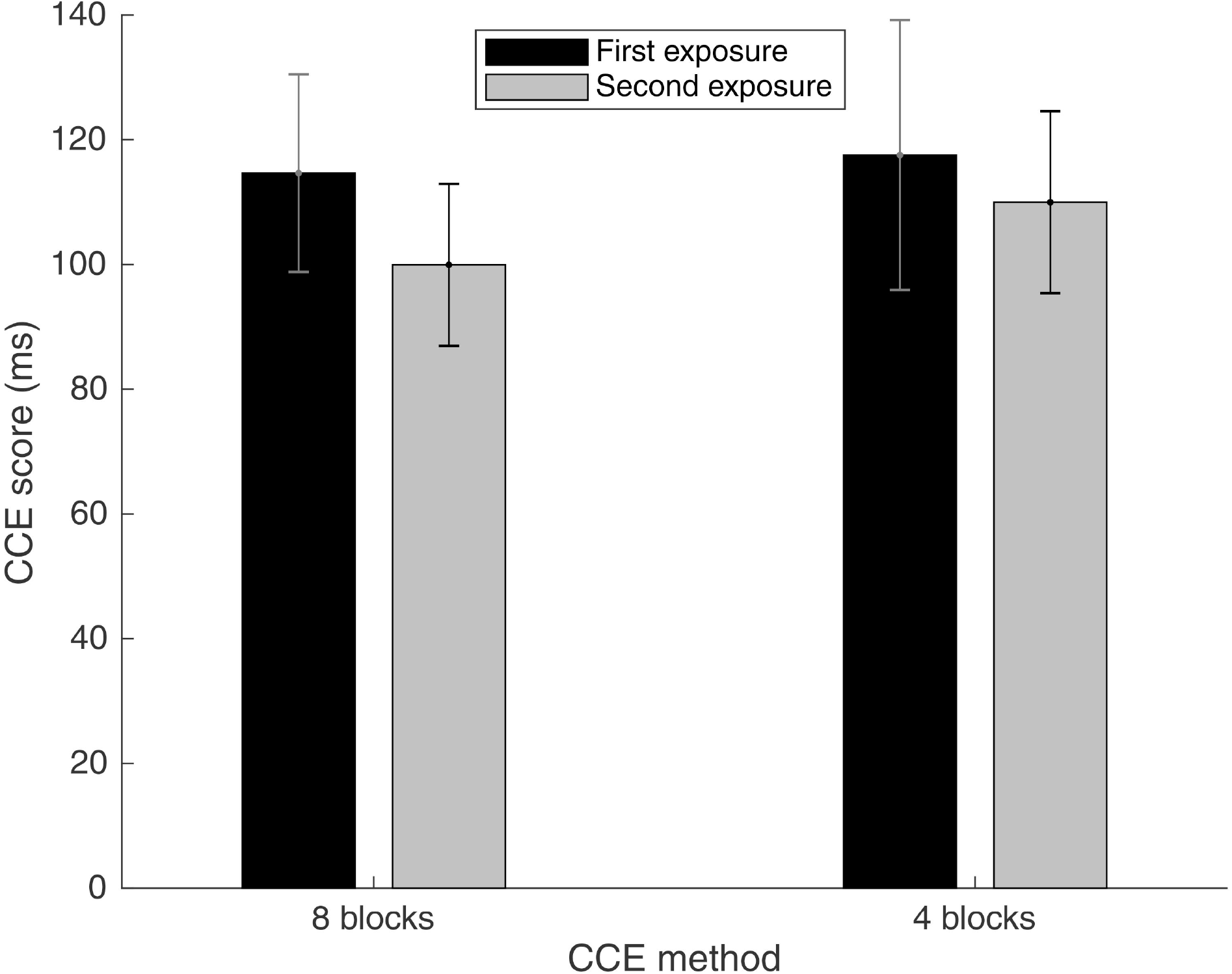
First exposure vs. second exposure mean CCE scores for 8-block and 4-block protocols. Means and standard error for 8-block (n=9) and 4-block (n=9) participants. Dark colored bars represent the first exposure session and light colored bars represent the second exposure session.

## Discussion

The CCE is a well-established quantitative measure of feedback incorporation that has important implications for objective human sensorimotor assessments (Spence et al., 1998; Spence, Pavani & Driver 2004; Spence, Pavani, Maravita & Holmes 2004; Spence 2015). The resulting scores have been used to quantify the rubber hand illusion (Zopf et al., 2010) and show promise for assessing neuroprosthetic devices (Spence 2015). Many applications of the CCE task require repeated testing – for example, testing the same subject under different sensory feedback conditions to see which one enables the highest degree of feedback incorporation. Oftentimes in research or clinical settings time allotted for sensorimotor assessment can be limited. This study sought to characterize the CCE task performance through repeat testing to inform its future implementation.

In Experiment 1, we found that with repeat testing, CCE scores decreased over the initial exposure sessions. The results are not explained by a fatigue effect as the CCE score trend was not matched by the overall reaction times measured with a ruler grasp task. Different motivation levels or strategies fail to explain the CCE score trend as CCE error rate did not show a trend matching the decreasing CCE scores. Thus, we consider the decrease in CCE score to be evidence of a task learning effect.

In Experiment 2, we demonstrated that the learning effect is persistent as CCE scores after six months remained near the stabilized scores observed at the end of five days of consecutive testing. The persistence of the learning effect provides an opportunity for researchers to only use CCE scores collected after learning has stabilized.

In certain circumstances, additional testing to arrive at stable CCE scores may not be feasible. In Experiment 3, we showed that a reduced-duration CCE protocol with 4 testing blocks instead of 8 reduced the impact of the learning effect across two testing sessions. Importantly, CCE score variability did not significantly increase with the 4-block protocol.

Reducing the number of blocks per session has the added benefit of reducing fatigue, improving the ability of subjects to concentrate over the duration of the study, and shortening the length of the session to approximately 30 minutes (which in turn makes it viable to test a greater number of subjects or for use in a clinical setting). Using a modified version of the CCE with only four blocks of 64 trials has many potential benefits, with the only drawback identified being a slight increase in the variability of results.

Through this characterization of the learning effect associated with the CCE task, we have provided support for two distinct mitigation strategies. If extended testing is feasible, researchers can provide enough practice so that CCE scores are only measured and compared after task learning has stabilized. In situations where reduced testing is required, researchers can use the 4-block CCE protocol to reduce overall task exposure and subsequently reduce the impact of the learning effect on the results. It is important that these strategies not be mixed; CCE scores should only be compared to other scores that were collected in the same manner.

It is not obvious what is being ‘learned’ through repeated task exposure. We suspect that subjects strengthen an internal model of the foot pedal location tied to the vibration stimulus, resulting in faster response times. Anecdotal support for this idea comes from one subject’s comment that “When the feedback is in the opposite location to the light, I sometimes find myself thinking about which pedal to press only to realize I’ve already pressed the correct pedal”. Somatosensory reorganization of an individual’s body schema through training may also be implicated (Cardinali et al., 2009).

The initial session CCE scores (i.e. before learning effect stabilization) appear to be higher than scores reported by other researchers (Spence, Pavani & Driver 2004; Zopf, Savage & Williams 2010). One potential explanation for this observation is that other studies may have different baseline CCE scores due to the experimental inquiry, such as studies using spatial misalignment (Spence, Pavani & Driver 2004) or artificial hands (Zopf, Savage & Williams 2010; Marini et al., 2014). We would expect a lower degree of feedback incorporation, and thus a lower CCE score (Maravita et al. 2002), under these conditions compared to the ideal conditions we used with both vibratory feedback and visual distractors aligned in place on one of the subject’s hands. Additionally, different experimental setups could account for differences in CCE scores. Others have recorded user inputs differently, e.g with a rocking heel-toe foot pedal setup (Spence, Pavani & Driver 2004) or with finger presses (Zopf, Savage & Williams 2010). Other variable techniques include the exclusion of trials with extensive eye movement (over 9% omitted in one study (Spence, Pavani & Driver 2004)), or the use of no-go trials as a control (Zopf, Savage & Williams 2010).

Another possible explanation for the observed variability in CCE scores across studies is the different methods used to apply feedback. Different feedback modalities result in different CCE scores (Frigs & Spence 2010; Mayer, Franco, Canive & Harrington 2009). Even when comparing studies using vibration feedback, the method of application can vary and may result in CCE score differences. We used small vibratory motors (0.8cm diameter vibrating surface) but others have used larger bone conduction vibrators (1.6cm x 2.4cm vibrating surface, (Spence, Pavani & Driver 2004)) or small speakers (0.9cm diameter vibrating surface, (Zopf, Savage & Williams 2010)), all could result in different degrees of feedback incorporation.

Differences in subject characteristics could also explain differences in CCE scores. All of the subjects in this study were initially naïve to the CCE task, but it is unclear if that was an inclusion criteria in other studies. Even comparing the first session results from this study across the two cohorts of subjects tested with 8 CCE test blocks we see differences in mean CCE. In Experiment 1, a mean CCE score of 140ms was observed (Fig 2) compared to 115ms in Experiment 3 (Fig 5). To summarize, there are lots of factors that could affect CCE score and it is impossible to determine if results reported elsewhere are capturing pre-, mid-or post-learning effect scores.

To our knowledge, this is the first study that has demonstrated the modulation of CCE score due to task overexposure. We recognize some limitations with this study that could warrant additional studies. Although we collected a generalized reaction time metric with a ruler grasp task, more sophisticated tracking of mental and physical fatigue would be helpful to better inform our understanding of the observed learning effect. Although CCE testing for individual participants was conducted at the same time of the day for each exposure, no strict guidelines were used to ensure that participants were in similar mental or physical states before each test session. Nevertheless, the learning effect we have described has important implications for future use of the CCE task. Going forward, the learning effect must be considered when measuring CCE scores by only using scores from after learning has stabilized, or by using a modified protocol that reduces exposure to the task.

## References

Cardinali, L., Frassinetti, F., Brozzoli, C., Urquizar, C., Roy, A.C., & Farnè, A. (2009). Tool-use induces morphological updating of the body schema. Current Biology, 19(12), R478–9. doi:10.1016/j.cub.2009.05.009

Frings, C., & Spence, C. (2010). Crossmodal congruency effects based on stimulus identity. Brain Research, 1354, 113–22. doi:10.1016/j.brainres.2010.07.058

Holmes, N.P., Calvert, G.A., & Spence, C. (2007). Tool use changes multisensory interactions in seconds: evidence from the crossmodal congruency task. Experimental Brain Research, 183(4), 465–76. doi:10.1007/s00221-007-1060-7

Holmes, N.P., & Spence, C. (2004). The body schema and the multisensory representation(s) of peripersonal space. Cognitive Processing, 5(2), 94–105. doi:10.1007/s10339-004-0013-3

Maravita, A., Spence, C., & Driver, J. (2003). Multisensory integration and the body schema: close to hand and within reach. Current Biology, 13(13), R531–9. doi:10.1016/S0960-9822(03)00449-4

Maravita, A., Spence, C., Kennett, S., & Driver, J. (2002). Tool-use changes multimodal spatial interactions between vision and touch in normal humans. Cognition, 83(2), B25–34. doi:10.1016/S0010-0277(02)00003-3

Marini, F., Tagliabue, C.F., Sposito, A.V., Hernandez-Arieta, A., Brugger, P., Estévez, N., Maravita, A. (2014). Crossmodal representation of a functional robotic hand arises after extensive training in healthy participants. Neuropsychologia, 53, 178–86. doi:10.1016/j.neuropsychologia.2013.11.017

Mayer, A.R., Franco, A.R., Canive, J., & Harrington, D.L.. (2009). The effects of stimulus modality and frequency of stimulus presentation on cross-modal distraction. Cerebral Cortex, 19(5), 993–1007. doi:10.1093/cercor/bhn148

Poliakoff, E., Ashworth, S., Lowe, C., & Spence, C. (2006). Vision and touch in ageing: crossmodal selective attention and visuotactile spatial interactions. Neuropsychologia, 44(4), 507–17. doi:10.1016/j.neuropsychologia.2005.07.004

Sengül, A., Rognini, G., van Elk, M., Aspell, J.E., Bleuler, H., Blanke, O. (2013). Force feedback facilitates multisensory integration during robotic tool use. Experimental Brain Research, 227(4), 497–507. doi:10.1007/s00221-013-3526-0

Sengül, A., van Elk, M., Rognini, G., Aspell, J.E., Bleuler, H., & Blanke, O. (2012). Extending the body to virtual tools using a robotic surgical interface: evidence from the crossmodal congruency task. PLoS ONE, 7(12), e49473. doi:10.1371/journal.pone.0049473

Spence, C. (2015). The Cognitive Neuroscience of Incorporation: Body Image Adjustment and Neuroprosthetics. In: K. Kansaku, L. Cohen, & N. Birbaumer (Eds.), Clinical Systems Neuroscience (pp. 151–68). Tokyo: Springer. doi:10.1007/978-4-431-55037-2_9

Spence, C., Kingstone, A., Shore, D.I., & Gazzaniga, M.S. (2001). Representation of visuotactile space in the split brain. Psychological Science, 12(1), 90–3. doi:10.1111/1467-9280.00316

Spence, C., Nicholls, M. E., Gillespie, N., & Driver, J. (1998). Cross-modal links in exogenous covert spatial orienting between touch, audition, and vision. Perception & Psychophysics, 60(4),544–57. doi:10.3758/BF03206045

Spence, C., Pavani, F., & Driver, J. (2004). Spatial constraints on visual-tactile cross-modal distractor congruency effects. Cognitive, Affective, & Behavorial Neuroscience, 4(2), 148–69. doi:10.3758/CABN.4.2.148

Spence, C., Pavani, F., Maravita, A., & Holmes, N. (2004). Multisensory contributions to the 3-D representation of visuotactile peripersonal space in humans: evidence from the crossmodal congruency task. Journal of Physiology - Paris, 98(1-3), 171–89. doi:10.1016/j.jphysparis.2004.03.008

Weir, J.P. (2005). Quantifying test-retest reliability using the intraclass correlation coefficient and the SEM. Journal of Strength & Conditioning Research, 19(1), 231–40. doi:10.1519/15184.1

Zopf, R., Savage, G., & Williams, M.A. (2010). Crossmodal congruency measures of lateral distance effects on the rubber hand illusion. Neuropsychologia, 48(3), 713–25. doi:10.1016/j.neuropsychologia.2009.10.028

Zopf, R., Savage, G., & Williams, M.A. (2013). The Crossmodal Congruency Task as a Means to Obtain an Objective Behavioral Measure in the Rubber Hand Illusion Paradigm. Journal of Visualized Experiments, (77), e50530. doi:10.3791/50530

